# Pre-training with pseudo-labeling compares favorably with large language models for regulatory sequence prediction

**DOI:** 10.1101/2023.12.21.572780

**Authors:** Raphaël Mourad

## Abstract

Predicting molecular processes using deep learning is a promising approach to provide biological insights for non-coding SNPs identified in genome-wide association studies. However, most deep learning methods rely on supervised learning, which requires DNA sequences associated with functional data, and whose amount is severely limited by the finite size of the human genome. Conversely, the amount of mammalian DNA sequences is growing exponentially due to ongoing large-scale sequencing projects, but in most cases without functional data. To alleviate the limitations of supervised learning, we propose a novel semi-supervised learning based on pseudo-labeling, which allows to exploit unlabeled DNA sequences from numerous genomes during model pre-training. The approach is very flexible and can be used to train any neural architecture including state-of-the-art models, and shows in certain situations strong predictive performance improvements compared to standard supervised learning in most cases. Moreover, small models trained by SSL showed similar or better performance than large language model DNABERT2.

## 1 Introduction

Complex genetic diseases are pathologies that are caused by a set of mutations, lifestyle and environmental factors [1]. Those diseases are highly prevalent in the population and include some cancers, heart diseases, neurological disorders and autoimmune diseases [2–4]. Over the past 20 years, genome-wide association studies (GWASs) have comprehensively mapped thousands of single nucleotide polymorphisms (SNPs) associated with these diseases [1]. However, the majority of those SNP were found outside coding sequences, make it difficult to gain insight into the underlying biological pathways [5].

Deep neural networks are increasingly used for regulatory element prediction from the DNA sequence. Convolutional neural networks (CNNs) were initially proposed, and achieved very good performances [6,7]. Nevertheless, CNNs could not capture long-range interactions between distant DNA motifs. To tackle this issue, other models were proposed instead to model long-range interactions, including recurrent neural networks with LSTM for few hundred bases [8] and dilated convolutions [9] for hundreds of kilobases. Those models were classically trained with labeled data only (supervised learning), which are limited by the finite size of the human genome (3.3 Gb).

Self-supervised learning with large language models (transformers) were recently trained on DNA sequences without labeled data, as a pretraining step before a fine-tuning step [10–16]. While such approach showed great performances, the training of large models (*>*100M parameters) on a large quantity of data (several genomes) requires important computational ressources, *i*.*e*. a cluster of GPUs or TPUs. An alternative to self-supervised learning is semi-supervised learning which uses during training a mix of labeled and unlabeled data [17]. Unlike other natural languages, DNA sequences are texts that are phylogenetically conserved among species due to the evolution process. In particular, it is known that many regulatory sequences and their associated functions are strongly conserved between close species. Therefore, conservation of regulatory sequences between closely related genomes could be used to greatly increase the size of the data by pseudo-labeling of sequences.

Here, we propose a novel semi-supervised learning (SSL) method based on cross-species pseudo-labeling, which greatly augments the size of the available labeled data for learning. The method consists in remapping regulatory sequences from a labeled genome (*e*.*g*. human genome) to other closely related genomes (*e*.*g*. mammalian genomes). Pseudo-labeled data allows to pretrain a neural network from multiple orders of magnitude larger data than the original labeled data. After pretraing with pseudo-labeled data, the model is then fine-tuned on the labeled data. The proposed SSL was used to train multiple state-of-the-art models, including DeepBind, DeepSea and DNABERT2, and showed sequence classification accuracy improvement in many cases. We also found an improvement in SNP effect prediction with SSL, especially for specific TFs.

## 2 Materials and Methods

### 2.1 Human experimental data

We used publicly available ATF3, ETS1, REST, MAX, P300, RAD21, CTCF, H3K4me3, POL2 and ANDR ChIP-seq data and input data of human lymphoblastoid GM12878 from Gene Expression Omnibus (GEO) accession GSE31477, GSE170139, GSE104399, GSE95899 from ENCODE [18]. We used publicly available ATAC-seq data of lymphoblastoid GM12878 from Gene Expression Omnibus (GEO) accession GSE170918 from ENCODE [18]. All the data were mapped on hg38 and peak calling was done using macs3 [19].

### 2.2 Human data labeling

#### 2.2.1 Classification for benchmarking with DeepBind and DNABERT2

We binned the human genome (*e*.*g*. human genome assembly hg38) into non-overlapping genomic intervals of 200 b, where a given bin was considered as either bound by a TF (if overlapping *>* 50% a peak), or not bound otherwise. For each 200 b bin, we extracted the corresponding DNA sequence (only containing the bin).

#### 2.2.2 Classification for benchmarking with DeepSea

We used the same approach as in DeepSea [7], described as follows. We binned the human genome (*e*.*g*. human genome assembly hg38) into non-overlapping genomic intervals of 200 b, where a given bin was considered as either bound by a TF (if overlapping *>* 50% a peak), or not bound otherwise. For each 200 b bin, we extracted the surrounding DNA sequence of 1 kb (containing the bin and the context).

### 2.3 Cross-species data labeling

To search for homologous sequences, we used the following 22 mammalian genomes: Rhesus monkey (rheMac10), marmoset (calJac3), chimpanzee (panTro6), pygmy chimpanzee (panPan2), Sumatran orangutan (ponAbe3), gorilla (gorGor6), olive baboon (papAnu4), crab-eating macaque (macFas5), Bolivian squirrel monkey (saiBol1), northern white-cheeked gibbon (nomLeu3), gray mouse lemur (micMur2), small-eared galago (otoGar3), mouse (mm10), rat (rn7), ferret (musFur1), rabbit (oryCun2), pork (susScr11), cat (felCat9), dog (canFam3), horse (equCab3), cow (bosTau9) and opossum (monDom5).

For pseudo-labeling, we liftovered peaks (ChIP-seq, ATAC-seq) from the human genome (labeled genome) to unlabeled genomes as follows. The UCSC liftover program was used to liftover the peak coordinates from the labeled genome to the unlabeled genomes. After liftover, a peak could be split into a set of non-overlapping regions. If a peak mapped to different loci in another genome, but the different loci remained close to each other (separated by *<* 20 b), the different loci were merged into one loci (homologous peak). Then, the corresponding DNA sequence was extracted and considered as a homologous sequence. If the peak mapped to distant loci (separated by *>*= 20 b) or if the peak did not map to any loci, then no homologous peak was found and no homologous sequence was extracted.

### 2.4 Models

#### 2.4.1 Shallow CNN

We have designed a simple shallow CNN (similar to DeepBind [6]) which consisted in a convolutional layer (64 or 256 filters and kernel size=24, activation=“relu”), a 1D global max pooling layer, a dropout layer (dropout rate=0.2), a dense layer (10 units), a dropout layer (dropout rate=0.2) and a last classification layer (1 unit, activation=“sigmoid”).

#### 2.4.2 Deep CNN

We have also designed a more sophisticated CNN (similar to Basenji [9]) which consisted in a convolutional layer (64 or 256 filters and kernel size=24, activation=“relu”), followed by 5 successive dilated convolutional layers with residual connection (64 or 256 filters and kernel size=3, activation=“relu”), a 1D global max pooling layer, a dropout layer (dropout rate=0.2), a dense layer (10 units), a dropout layer (dropout rate=0.2) and a last classification layer (1 unit, activation=“sigmoid”).

#### 2.4.3 DeepBind

We reimplemented the DeepBind model using Keras, keeping the original model architecture [6], a convolutional layer (16 filters and kernel size=24, activation=“relu”), a 1D global max pooling layer, a dense layer (32 units), a dropout layer (dropout rate=0.5) and a last classification layer (1 unit, activation=“sigmoid”).

#### 2.4.4 DeepSea

We reimplemented the DeepSea model using Keras keeping the original model architecture [7]: a convolutional layer (320 filters and kernel size=8, activation=“relu”), a 1D max pooling layer (pool size=4, strides=4), a dropout layer (dropout rate=0.2), a convolutional layer (480 filters and kernel size=8, activation=“relu”), a 1D max pooling layer (pool size=4, strides=4), a dropout layer (dropout rate=0.2), a convolutional layer (960 filters and kernel size=8, activation=“relu”), a dropout layer (dropout rate=0.5), a flatten layer, a dense layer (925 units) and a last classification layer (1 unit, activation=“sigmoid”).

#### 2.4.5 DNABERT2

We fine-tuned DNABERT2 [12] (downloaded from zhihan1996/DNABERT-2-117M huggingface repository) for classification with the following default parameters as provided with the “config.json” file: alibi_starting_size=512, architectures=“BertForMaskedLM”, attention_probs_dropout_prob=0.0, gradient_checkpointing=false, hidden_act=“gelu”, hidden_dropout_prob=0.1, hidden_size=768, initializer_range=0.02, intermediate_size=3072, layer_norm_eps=1e-12, max_position_embeddings=512, model_type=“bert”, num_attention_heads=12, num_hidden_layers=12, position_embedding_type=“absolute”, torch_dtype=“float32”, transformers_version=4.28.0, type_vocab_size=2, use_cache=true, vocab_size=4096.

### 2.5 Prediction of SNP effect

We used the previous deep learning models to predict the effect of SNPs on functional data, such as protein binding or chromatin accessibility. To compute the effect of a SNP, we used the following approach for classification as in [6]. First, model predictions were computed both for the DNA sequence comprising the reference SNP allele (*p*_*ref*_) and the same DNA sequence but with the alternative SNP allele (*p*_*alt*_). Then, the SNP effect was computed as (*p*_*alt*_ *− p*_*ref*_) *· max*(0, *p*_*alt*_, *p*_*ref*_), where *p*_*alt*_ is the prediction from the alternative allele and *p*_*ref*_ the prediction from the reference allele.

### 2.6 Implementation and Availability

The model was developed using Tensorflow and Keras. It is available at the Github repository: https://forgemia.inra.fr/raphael.mourad/deepssl

## 3 Results and Discussion

### 3.1 Cross-species data labeling approach and novel training strategy

The proposed cross-species pseudo-labeling approach is illustrated in Figure 1A. Given a labeled sequence (*e*.*g*. ChIP-seq peak in yellow), cross-species data labeling consists in identifying the homologous sequences in related species (*e*.*g*. mammals) using liftover from the labeled genome (*e*.*g*. the human genome) to the unlabeled genomes (*e*.*g*. the chimpanzee, the dog and the rabbit genomes). In Figure 1A, a homologous sequence was found in the chimpanzee (in orange) and in the dog (in blue), but no homologous sequence was found in the rabbit. Cross-species labeled data can then be used to “augment” the original labeled data (*e*.*g*. the human genome), in sense that it can considerably increase the amount of labeled data that could be used for pre-training the model.

**Figure 1.**
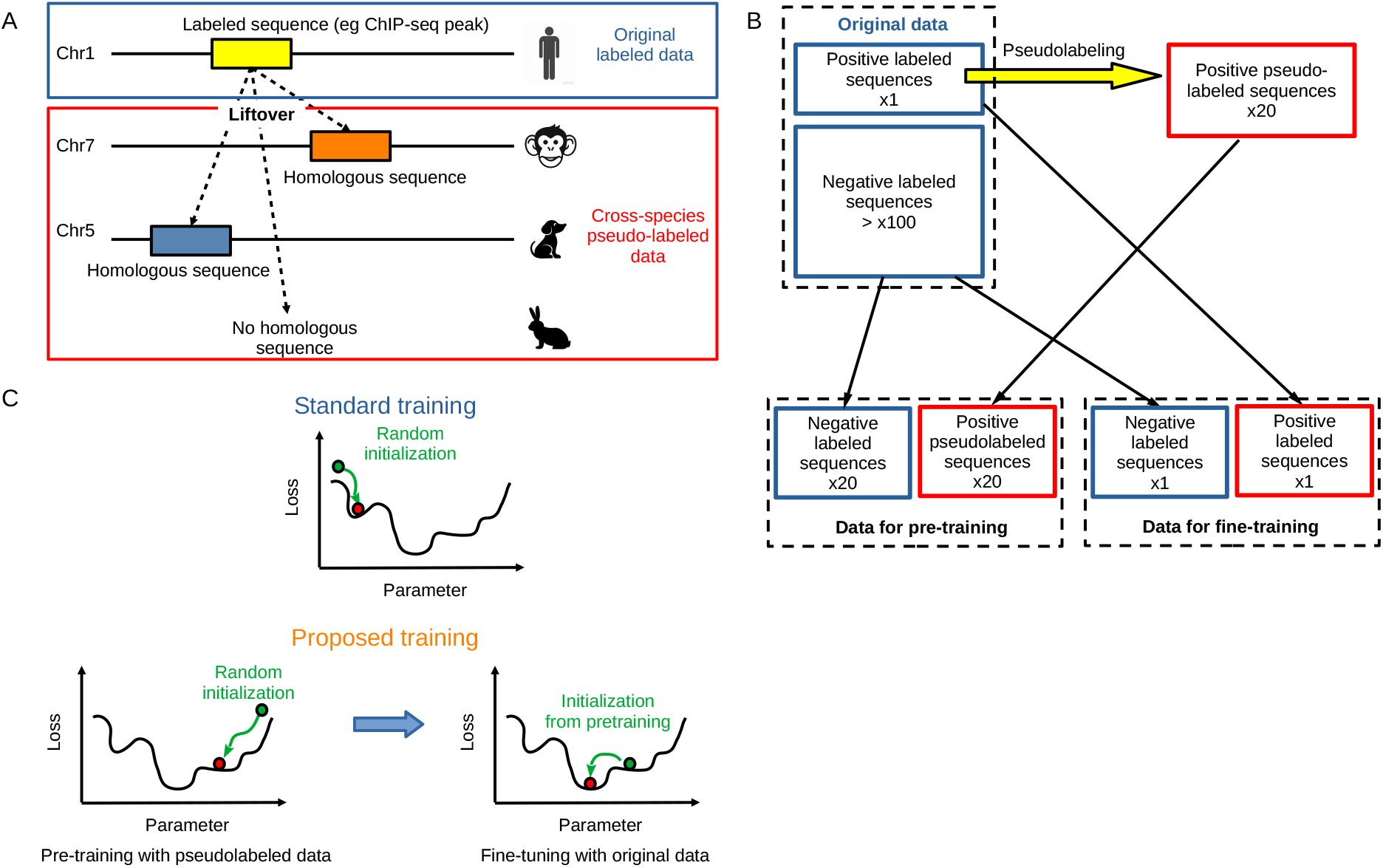
Data pseudo-labeling and associated semi-supervised learning. A) Data pseudo-labeling using cross-species labeling by homology. B) Data used for pre-training and for fine-tuning. C) Semi-supervised learning based on pre-training with pseudo-labeled data, followed by fine-tuning with the original labeled data.

Using the cross-species pseudo-labeled data, we designed a novel strategy to train more efficiently deep learning models. The strategy consists in two steps:

- first, pre-training of the model using cross-species pseudo-labeled data. For this purpose, the model is trained with balanced data comprising positive sequences corresponding to all sequences pseudo-labeled as overlapping a peak (*i*.*e*. non-human sequences that are homologous to human sequences overlapping a peak), and the same number of negative sequences corresponding to sequences labeled as non-overlapping a peak (*i*.*e*. human sequences) (Figure 1B).
- second, fine-tuning of the model using the original labeled data (*e*.*g*. human labeled sequences). The model is trained with balanced data comprising positive sequences corresponding to sequences labeled as overlapping a peak (human sequences), and the same number of negative sequences corresponding to sequences labeled as non-overlapping a peak (*i*.*e*. human sequences) (Figure 1B).

The rationale behind is that the pre-training step with pseudo-labeled data will provide a better parameter initialization for the fine-tuning with the original labeled data (Figure 1C).

### 3.2 Improving state-of-the-art model performance with semi-supervised learning

We then applied our novel training scheme with three state-of-the-art models, namely: DeepBind (shallow CNN), DeepSea (deep CNN) and DNABERT2 (large language model). For each model, we compared prediction performances for sequence classification for different experimental data between standard supervised learning (*i*.*e*. without pretraining on pseudo-labeled data) and our novel semi-supervised learning (*i*.*e*. with pretraining on pseudo-labeled data).

For a comprehensive benchmarking, we ran the comparison on diverse experiments:

- ChIP-seq of ATF3, a specific TF binding to 1677 loci with a median peak size of around 170 pb;
- ChIP-seq of ETS1, a specific TF binding to 4120 loci with a median peak size of around 224 pb;
- ChIP-seq of REST, a specific TF binding to 6119 loci with a median peak size of around 276 pb;
- ChIP-seq of MAX, a specific TF binding to 13605 loci with a median peak size of around 421 pb;
- ChIP-seq of P300, a specific TF binding to 14223 loci with a median peak size of around 217 pb;
- ChIP-seq of RAD21, a wide-spread TF binding to 34623 loci with a median peak size of around 265 pb;
- ChIP-seq of CTCF, a wide-spread TF binding to 72779 loci with a median peak size of around 210 pb;
- ChIP-seq of H3K4me3, a histone modification that marks promoters and that is present at 25641 loci with a median peak size of around 438 pb;
- ChIP-seq of POL2, a wide-spread multiprotein complex that transcribes DNA into pre-mRNA and that binds to 35982 loci with a median peak size of around 250 pb;
- ATAC-seq, a non specific assay which maps accessible chromatin and which was observed at 102030 loci (or peaks) with a median peak size of around 230 pb;

The benchmarking was done in a classification setting (presence/absence of a peak in a bin). Models were trained on all chromosomes, except chromosomes 8 and 9 that were kept for testing. In Table 1, for each data, we show the number of positive sequences and the number of pseudo-labeled positive sequences obtained by pseudo-labeling on 22 mammalian genomes. For instance, for ATF3, a TF that binds to very few sequences (1306), the number of positive sequences obtained by pseudo-labeling was 23555, which corresponds to an increase by *×*18. Of note, pseudo-labeling with 22 mammalian genomes does not increase by *×*22 the amount of labeled sequences, as there are many sequences that are not evolutionary conserved. However, such approach is very efficient, as in average 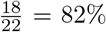 of sequences are conserved and therefore could be pseudo-labeled.

**Table 1.** Number of positive sequences and pseudo-labeled positive sequences obtained by pseudo-labeling with 22 mammalian genomes for each data.

| Data | Number of positive sequences | Number of pseudo-labeled Positive sequences | Increase |
| --- | --- | --- | --- |
| ATF3 | 1306 | 23555 | 18x |
| ETS1 | 4211 | 77661 | 18x |
| REST | 7201 | 130615 | 18x |
| MAX | 23916 | 441295 | 18x |
| P300 | 15844 | 272923 | 17x |
| RAD21 | 37113 | 646495 | 17x |
| CTCF | 75082 | 1245195 | 17x |
| H3K4me3 | 67589 | 1220489 | 18x |
| POL2 | 57226 | 998107 | 17x |
| ATAC | 70600 | 1186950 | 17x |

#### 3.2.1 Implementation with DeepBind and with DeepSea (CNNs)

We first comprehensively compared classical supervised learning (SL) with our novel SSL training for DeepBind (Figure 2A). In term of area under the receiver operating characteristic (AUROC) curve, we observed an overall significant increase using SSL (p=2 *×* 10^*−*9^). There was a large increase with SSL compared to SL for ATF3 (+11.3%), REST (+7.4%) and P300 (+24.4%). But for other experiments, gains were slight, around 1 *−* 2%. However, as the data were highly inbalanced with more negative sequences, the area under the precision recall (AUPR) curve was more appropriate. In term of AUPR, improvements were larger for most experiments (overall significance p=1 *×* 10^*−*9^). For instance, there was a much higher increase for RAD21 (+48%), CTCF (+36%) or POL2 (+29.7%). For TFs with few binding sites (specific TFs), there was a very strong increase, as illustrated for ATF3 (+2323.8%), ETS1 (+133.6%) and REST (+794.2%).

**Figure 2.**
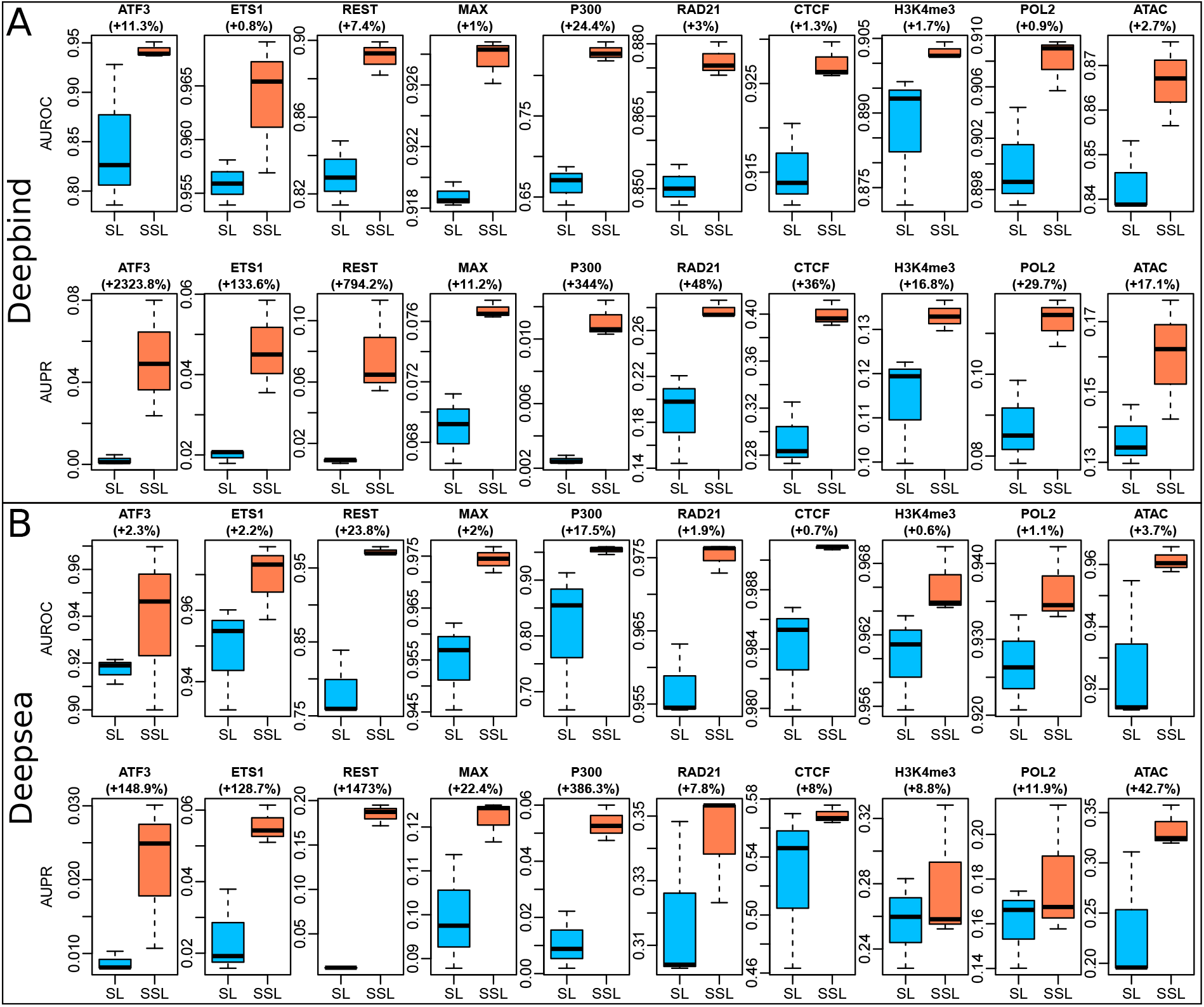
Benchmarking with DeepBind and DeepSea. AUROC: area under the receiver operating characteristics curve. AUPR: area under the precision recall curve. Each model was run 3 times.

We then compared SL with SSL for DeepSea (Figure 2B). In term of AUROC, we found a significant increase using SSL (p=3 *×* 10^*−*9^). A large increase was found with SSL compared to SL for REST (+23.8%) and P300 (+17.5%). But for other data, gains were slight, around 1 *−* 2%. In term of AUPR, as with DeepBind, improvements were larger for most experiments (p=6 *×* 10^*−*6^). For instance, there was a high increase for ATAC (+42.7%). As for DeepBind, larger increases were found for specific TFs, *e*.*g*. ATF3 (+148.9%), ETS1 (+128.7%) and REST (+1473%).

#### 3.2.2 Implementation with DNABERT2 (large language model)

Large language models (LLMs), in particular transformers, are currently the most powerful neural networks for natural language processing, and were recently adapted for DNA sequences. DNABERT2 is a transformer-based LLM which was previously pretrained by self-supervised learning on genomes from 135 species, comprising 32.49 billion bases. Pretrained DNABERT2 can then be fine-tuned on a given labeled data.

Here, we used pretrained DNABERT2 to be further trained by SL or by SSL. SL consisted in fine-tuning pretrained DNABERT2 on labeled data. SSL instead consisted in further pretraining of pretrained DNABERT2 with pseudo-labeled data and then fine-tuning on labeled data. Overall, there was a significant increase for AUROC (p= 0.002), but not for AUPR (p= 0.963). More specifically, we found that SSL strongly increased AUROCs and AUPRs for specific TFs ATF3 (AUROC: +2.2%, AUPR: +18.8%), ETS1 (AUROC: +4.2%, AUPR: +62.9%) and REST (AUROC: +10%, AUPR: +118%), whose data for fine-tuning are small (less than 10K peaks) (Figure 3). For P300, we also observed increased AUROCs and AUPRs (AUROC: +6.1%, AUPR: +8.5%), but for other data, we found a decrease in AUROC and AUPR values. Such results suggest that SSL is beneficial for pretrained LLMs when fine-tuning is done on small data, which is the case for specific TFs.

**Figure 3.**
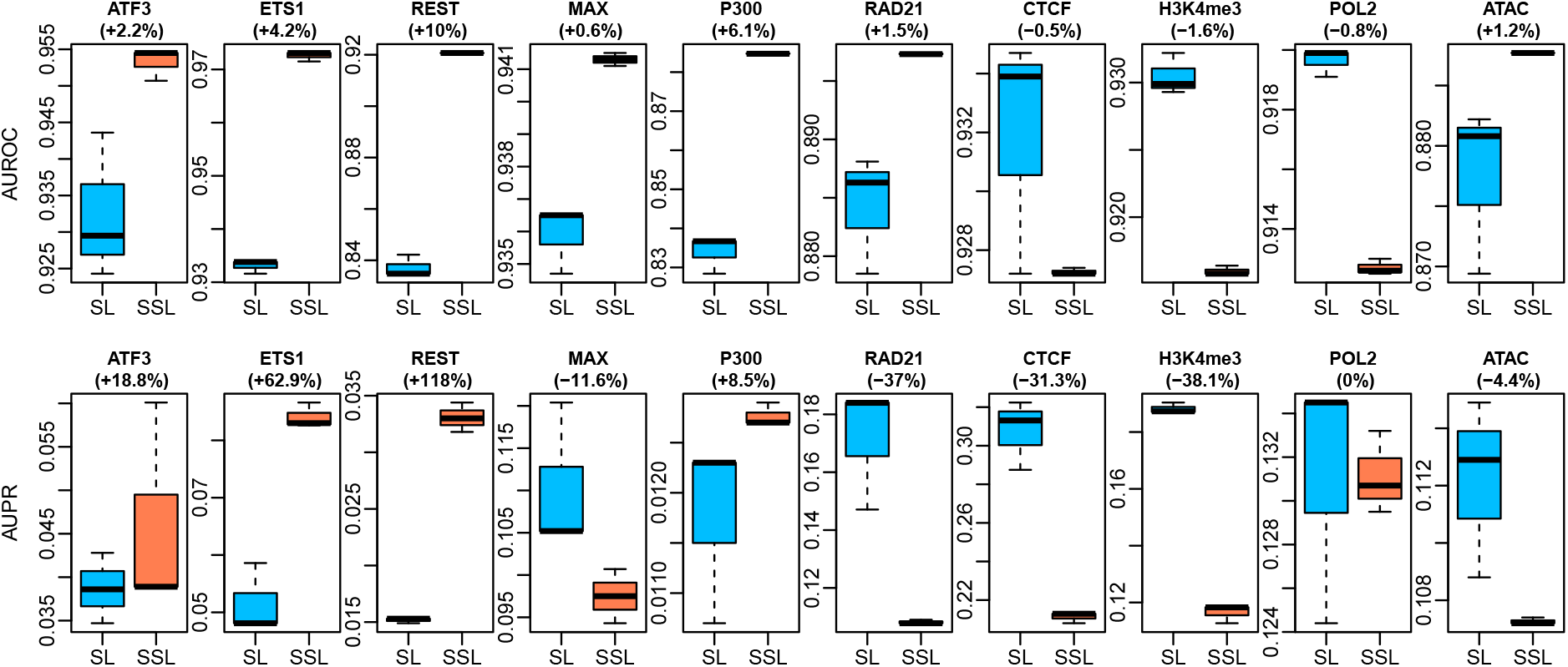
Benchmarking with DNABERT2. AUROC: area under the receiver operating characteristics curve. AUPR: area under the precision recall curve. Each model was run 3 times.

### 3.3 Small models trained with SSL versus large language model

In order to demonstrate the power of our new training approach, we then compared simple CNNs trained with SL or SSL, with state-of-the-art large language model DNABERT2 fine-tuned without SSL for different datasets. The simple CNNs only comprised 7K parameters (CNN-7K) or 27K parameters (CNN-27K), while DNABERT2 had around 117M parameters, respectively (therefore with a magnitude of order 3-4 compared to the CNNs).

Results are presented in Table 2. When comparing AUROCs, we found that the simple CNN-27K trained with SSL (CNN-27K-SSL) ranked first for 8 out of 10 data, compared to DNABERT2 which ranked first for 2 out of 10 data. For the AUPR which better accounts for class imbalance, CNN-27K-SSL also ranked first for 8 out of 10 data. We could see that AUPR improvements were huge with CNN-27K-SSL when compared to CNN-27K with SL (CNN-27K-SL), in particular for data with few peaks (small data). For instance, for ATF3, a TF that only binds to 1677 peaks, the AUPR was 0.149 for CNN-27K-SSL, which was 7 times higher than the AUPR for CNN-27K-SL (0.021). There was also an important increase for ETS1 with an AUPR of 0.139 for CNN-27K-SSL compared to 0.030 for CNN-27K-SL, and for REST with an AUPR of 0.253 for CNN-27K-SSL compared to 0.062 for CNN-27K-SL.

**Table 2.**
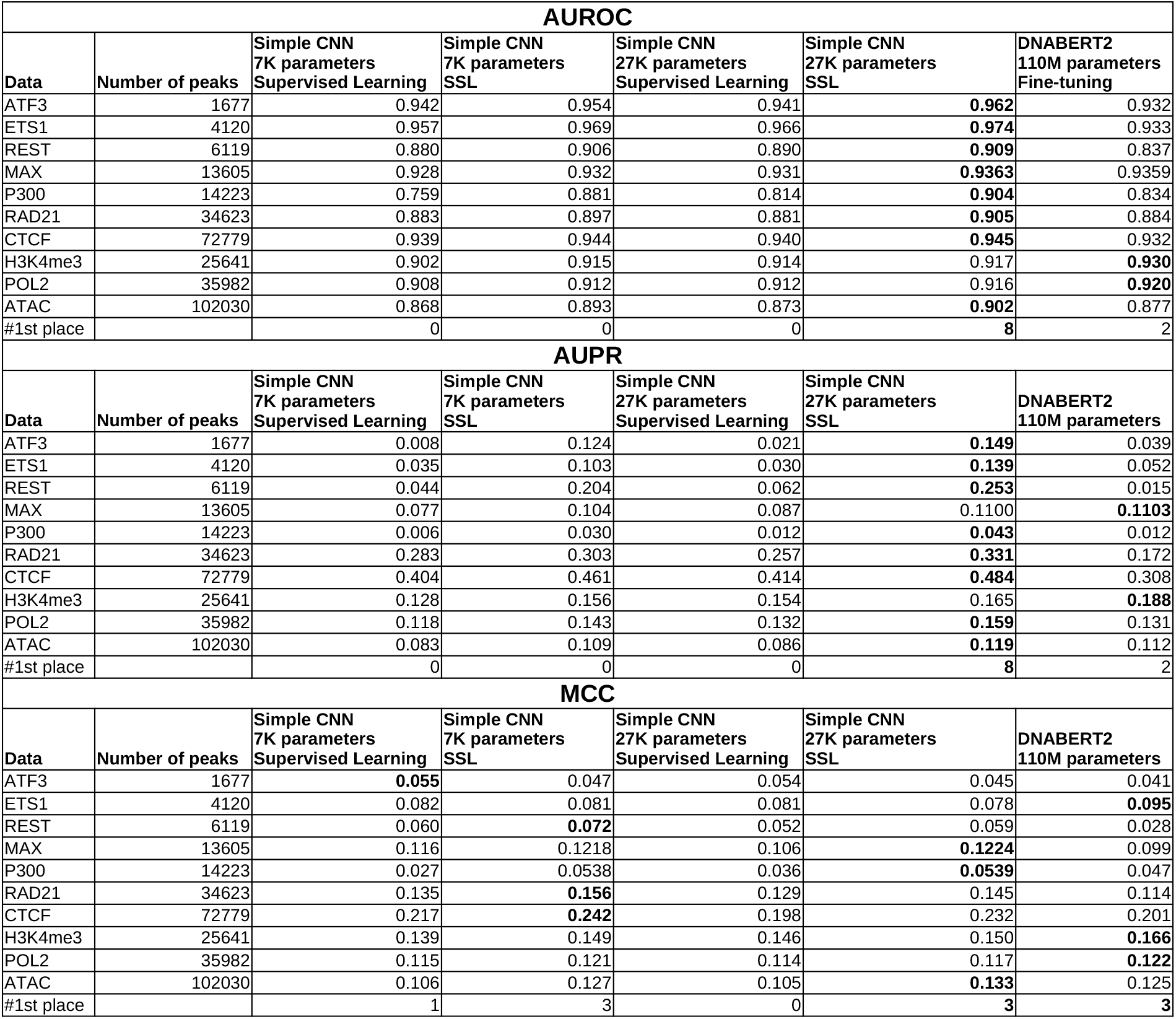
Benchmarking of a simple convolutional neural network (CNN) trained with SL or SSL, and comparison with large language model DNABERT2 on different datasets. AUROC: area under the receiver operating characteristics curve. AUPR: area under the precision recall curve. MCC: Matthew correlation coefficient. Each model was run 3 times and the average statistics were computed.

| AUROC |  |  |  |  |  |  |
| --- | --- | --- | --- | --- | --- | --- |
| Data | Number of peaks | Simple CNN<br>7K parameters<br>Supervised Learning | Simple CNN<br>7K parameters<br>SSL | Simple CNN<br>27K parameters<br>Supervised Learning | Simple CNN<br>27K parameters<br>SSL | DNABERT2<br>110M parameters<br>Fine-tuning |
| ATF3 | 1677 | 0.942 | 0.954 | 0.941 | **0.962** | 0.932 |
| ETS1 | 4120 | 0.957 | 0.969 | 0.966 | **0.974** | 0.933 |
| REST | 6119 | 0.880 | 0.906 | 0.890 | **0.909** | 0.837 |
| MAX | 13605 | 0.928 | 0.932 | 0.931 | **0.9363** | 0.9359 |
| P300 | 14223 | 0.759 | 0.881 | 0.814 | **0.904** | 0.834 |
| RAD21 | 34623 | 0.883 | 0.897 | 0.881 | **0.905** | 0.884 |
| CTCF | 72779 | 0.939 | 0.944 | 0.940 | **0.945** | 0.932 |
| H3K4me3 | 25641 | 0.902 | 0.915 | 0.914 | 0.917 | **0.930** |
| POL2 | 35982 | 0.908 | 0.912 | 0.912 | 0.916 | **0.920** |
| ATAC | 102030 | 0.868 | 0.893 | 0.873 | **0.902** | 0.877 |
| #1st place |  | 0 | 0 | 0 | **8** | 2 |
| AUPR |  |  |  |  |  |  |
| Data | Number of peaks | Simple CNN<br>7K parameters<br>Supervised Learning | Simple CNN<br>7K parameters<br>SSL | Simple CNN<br>27K parameters<br>Supervised Learning | Simple CNN<br>27K parameters<br>SSL | DNABERT2<br>110M parameters |
| ATF3 | 1677 | 0.008 | 0.124 | 0.021 | **0.149** | 0.039 |
| ETS1 | 4120 | 0.035 | 0.103 | 0.030 | **0.139** | 0.052 |
| REST | 6119 | 0.044 | 0.204 | 0.062 | **0.253** | 0.015 |
| MAX | 13605 | 0.077 | 0.104 | 0.087 | 0.1100 | **0.1103** |
| P300 | 14223 | 0.006 | 0.030 | 0.012 | **0.043** | 0.012 |
| RAD21 | 34623 | 0.283 | 0.303 | 0.257 | **0.331** | 0.172 |
| CTCF | 72779 | 0.404 | 0.461 | 0.414 | **0.484** | 0.308 |
| H3K4me3 | 25641 | 0.128 | 0.156 | 0.154 | 0.165 | **0.188** |
| POL2 | 35982 | 0.118 | 0.143 | 0.132 | **0.159** | 0.131 |
| ATAC | 102030 | 0.083 | 0.109 | 0.086 | **0.119** | 0.112 |
| #1st place |  | 0 | 0 | 0 | **8** | 2 |
| MCC |  |  |  |  |  |  |
| Data | Number of peaks | Simple CNN<br>7K parameters<br>Supervised Learning | Simple CNN<br>7K parameters<br>SSL | Simple CNN<br>27K parameters<br>Supervised Learning | Simple CNN<br>27K parameters<br>SSL | DNABERT2<br>110M parameters |
| ATF3 | 1677 | **0.055** | 0.047 | 0.054 | 0.045 | 0.041 |
| ETS1 | 4120 | 0.082 | 0.081 | 0.081 | 0.078 | **0.095** |
| REST | 6119 | 0.060 | **0.072** | 0.052 | 0.059 | 0.028 |
| MAX | 13605 | 0.116 | 0.1218 | 0.106 | **0.1224** | 0.099 |
| P300 | 14223 | 0.027 | 0.0538 | 0.036 | **0.0539** | 0.047 |
| RAD21 | 34623 | 0.135 | **0.156** | 0.129 | 0.145 | 0.114 |
| CTCF | 72779 | 0.217 | **0.242** | 0.198 | 0.232 | 0.201 |
| H3K4me3 | 25641 | 0.139 | 0.149 | 0.146 | 0.150 | **0.166** |
| POL2 | 35982 | 0.115 | 0.121 | 0.114 | 0.117 | **0.122** |
| ATAC | 102030 | 0.106 | 0.127 | 0.105 | **0.133** | 0.125 |
| #1st place |  | 1 | 3 | 0 | **3** | **3** |

When comparing Matthew correlation coefficients (MCCs) as used in [11], we found that CNN-7K-SSL, CNN-27K-SSL and DNABERT2 all ranked first for 3 out of 10 data.

Our benchmark showed that a simple CNN with only 27K parameters could outperform in certain situations or perform as well as LLM DNABERT2 which comprises millions of parameters and was trained on whole genomes. Moreover, the results revealed that our SSL strongly improved model performance for small data (*i*.*e*. specific TFs), which is expected as deep learning models comprise large numbers of parameters and therefore need large data for parameter learning.

### 3.4 Prediction of SNP effects

We next evaluated the ability of SSL to improve the prediction of SNP effect on molecular phenotypes such as TF binding, and compared it to SL. For this purpose, we first trained a shallow CNN with SL for CTCF peak classification, and then predicted the impact of a SNP on CTCF binding, as done in [7]. We found a good Spearman correlation between the observed effect as estimated by ChIP-seq allelic imbalance (from ADASTRA database [20]), and the predicted effect (R*s* = 0.383; Figure 4A). We then trained the same model with SSL and observed an increase of Spearman correlation (R*s* = 0.430, +12.3%; Figure 4A). We repeated the same experiment with a deep CNN with dilated convolutions and residual connections and obtained an R*s* = 0.419 for SL and an R*s* = 0.455 for SSL (+8.6%; Figure 4B). We changed the number of convolution kernels from 64 to 256. Results showed an increase of Rs with SSL for all model predictions (Figure 4C). We also trained CNNs to predict ANDR peaks and then predicted SNP effects. Compared to CTCF, which is a very wide-spread TF with 72779 loci, ANDR is a more specific TF with 3415 peaks, making the model harder to train with SL as fewer data are available. Interestingly, for ANDR, we found that the predictions of SNP effects were very bad (R*s <* 0) with SL, while there were positive correlations (R*s >* 0.25) with SSL for shallow CNNs and (R*s >* 0.15) for deep CNNs (Figure 4D).

**Figure 4.**
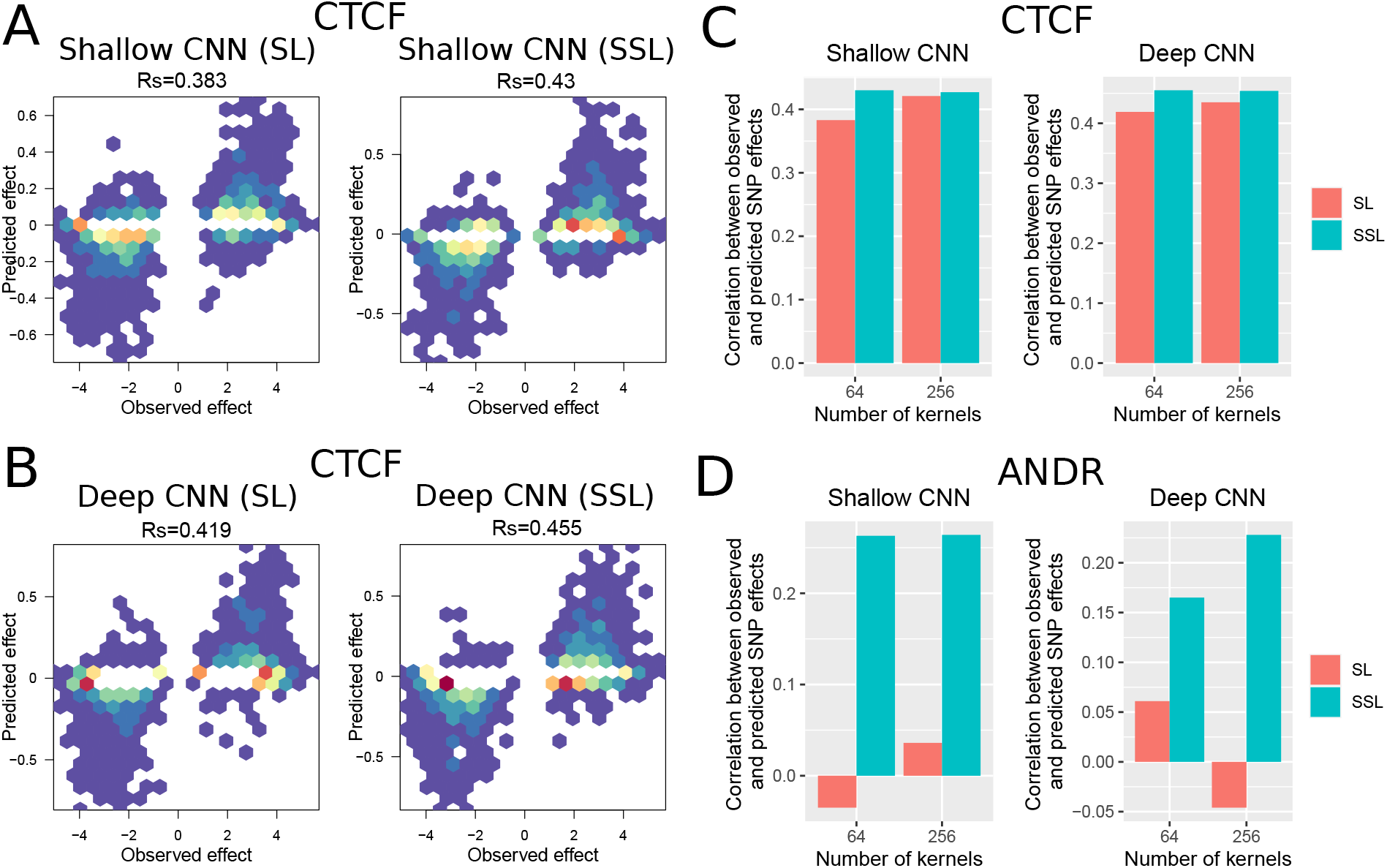
Prediction of SNP effects with supervised learning (SL) and semi-supervised learning (SSL). A) Scatter plots of observed CTCF SNP effects versus predicted effects for a shallow CNN trained with either SL or SSL. 64 kernels were used for the convolution. B) Same scatter plots for a deep CNN trained with either SL or SSL. 64 kernels were used for the convolution. C) Barplots of Spearman correlations between observed CTCF SNP effects versus predicted effects for shallow and deep CNNs, and 64 and 256 kernels. D) Same barplots for ANDR SNP effects.

Results thus showed that SSL can improve the prediction of SNP effects on molecular phenotypes,such as TF binding. Moreover, we found that the gain was much higher for specific TFs for which less data were available for model training.

## 4 Conclusion

In this article, we proposed a novel semi-supervised learning (SSL) based on pseudo-labeled data allowing to train models from data with multiple orders of magnitude compared to available labeled data. Such approach alleviates the size limit of the human genomes (3.3 Gb), by leveraging other mammmalian genomes. Given that the amount of sequenced mammmalian genomes is growing exponentially due to large-scale sequencing projets, such approach represents a promising avenue for improving the performance of current deep learning models.

Training of state-of-the-art models, including DeepBind, DeepSea and DNABERT2, showed that SSL outperformed SL for DNA sequence classification in many situations, in particular when training data were small, such as for specific TFs. Moreover, SSL showed higher performances compared to SL for predicting SNP effects, especially for specific TFs. In addition, benchmarking on a diverse dataset showed that a simple CNN of 27K parameters trained with SSL performs similarly or better than LLM DNABERT2 containing 110M parameters.

SSL is not the first approach that leverages genomes without functional data. Self-supervised learning with LLMs were recently trained on multiple genomes without labeled data, as a pretraining step before a fine-tuning step [11–16]. While such approach showed great performances, the training of large models (*>*100M parameters) on a large quantity of data (several complete genomes) requires important computational ressources, *i*.*e*. a cluster of GPUs or TPUs. SSL provides a complementary approach to pretrained LLMs by improving the accuracy of classical deep learning models (*e*.*g*. non-LLMs, for instance CNNs) in particular for specific TFs in term of AUPR. Compared to pretrained LLMs, SSL does not require to train large models with hundreds of millions of parameters. Moreover, by focusing only on homologous sequences from unlabeled genomes (*i*.*e*. only a small fraction of the genomes) and not the entire unlabeled genomes as LLMs, SSL trains models from more informative DNA sequences. Of note, the assets of SSL comes at the cost of task specificity, since the pretraining with pseudo-labeled data improves fine-tuning only for a given task. Conversely, pretrained LLMs have the main advantage to be fine-tuned on any task. Moreover, pretrained LLMs have zero-shot capabilities, *i*.*e*. for task for which the model hasn’t seen any data [21].

There are several limitations of the proposed approach. First, the pseudo-labeling assumes that the homogous sequence has the same label as the original sequence from which liftover was done. If the assumption does not hold, then SSL is expected to yield poor results, worst than those obtained with SL. To tackle this issue, during fine-tuning, the model can be used to discard homologous sequences with low predicted probability of the pseudo-label. Second, the use of our approach is likely limited to species which are not too evolutionary distant. For instance, it is very unlikely that including plant genomes would help to predict regulatory sequences from a mammalian genome. Third, the proposed SSL approach is very basic and more sophisticated methods can be used. For instance, the Noisy Student iterates between a teacher model that assigns labels to unlabeled data and a larger student model which adds noise to encourage the model’s decision making frontier to be smooth, on both labeled and unlabeled data [22]. Meta Pseudo Labels can also be used, where the teacher and the student models are trained in parallel, where the teacher learns to generate better pseudo labels and the student learns from the pseudo labels. [23].

## 5 Key Points

- Cross-species pseudo-labeling improves the training of deep learning models for regulatory sequence prediction by augmenting the data.
- Semi-supervised learning (SSL) is proposed to pretrain a model with pseudo-labeling data and then to fine-tune it on original labeled data.
- SSL shows strong performance increase for transcription factors with few binding sites (few labeled data) compared to classical supervised learning.
- A simple convolutional model (27K parameters) trained with SSL could outperform in certain situations or perform as well as large model DNABERT2 (117M parameters).

## Availability of data and materials

All the code and data are available at: https://forgemia.inra.fr/raphael.mourad/deepssl

## Authors’ contributions

RM conceived and designed the project. RM implemented the model and analyzed the data. RM wrote the manuscript.

## References

[1] Visscher, P. M., Wray, N. R., Zhang, Q., Sklar, P., McCarthy, M. I., Brown, M. A., and Yang, J. (July, 2017) 10 years of GWAS discovery: Biology, function, and translation. The American Journal of Human Genetics, 101(1), 5–22.

[2] Dorn, G. W. and Cresci, S. (February, 2009) Genome-wide association studies of coronary artery disease and heart failure: where are we going?. Pharmacogenomics, 10(2), 213–223 PMID: 19207022.

[3] Billings, L. K. and Florez, J. C. (November, 2010) The genetics of type 2 diabetes: What have we learned from GWAS?. Annals of the New York Academy of Sciences, 1212(1), 59–77.

[4] Collins, A. L. and Sullivan, P. F. (January, 2013) Genome-wide association studies in psychiatry: What have we learned?. The British Journal of Psychiatry, 202(1), 1–4.

[5] Maurano, M. T., Humbert, R., Rynes, E., Thurman, R. E., Haugen, E., Wang, H., Reynolds, A. P., Sandstrom, R., Qu, H., Brody, J., Shafer, A., Neri, F., Lee, K., Kutyavin, T., Stehling-Sun, S., Johnson, A. K., Canfield, T. K., Giste, E., Diegel, M., Bates, D., Hansen, R. S., Neph, S., Sabo, P. J., Heimfeld, S., Raubitschek, A., Ziegler, S., Cotsapas, C., Sotoodehnia, N., Glass, I., Sunyaev, S. R., Kaul, R., and Stamatoyannopoulos, J. A. (September, 2012) Systematic localization of common disease-associated variation in regulatory DNA. Science, 337(6099), 1190–1195.

[6] Alipanahi, B., Delong, A., Weirauch, M. T., and Frey, B. J. (July, 2015) Predicting the sequence specificities of DNA- and RNA-binding proteins by deep learning. Nature Biotechnology, 33, 831–838.

[7] Zhou, J. and Troyanskaya, O. G. (August, 2015) Predicting effects of noncoding variants with deep learning-based sequence model. Nature Methods, 12(10), 931–934.

[8] Quang, D. and Xie, X. (04, 2016) DanQ: a hybrid convolutional and recurrent deep neural network for quantifying the function of DNA sequences. Nucleic Acids Research, 44(11), e107–e107.

[9] Kelley, D. R., Reshef, Y. A., Bileschi, M., Belanger, D., McLean, C. Y., and Snoek, J. (2018) Sequential regulatory activity prediction across chromosomes with convolutional neural networks. Genome Research, 28(5), 739–750.

[10] Avsec, Ž., Agarwal, V., Visentin, D., Ledsam, J. R., Grabska-Barwinska, A., Taylor, K. R., Assael, Y., Jumper, J., Kohli, P., and Kelley, D. R. (Oct, 2021) Effective gene expression prediction from sequence by integrating long-range interactions. Nature Methods, 18(10), 1196–1203.

[11] Ji, Y., Zhou, Z., Liu, H., and Davuluri, R. V. (02, 2021) DNABERT: pre-trained Bidirectional Encoder Representations from Transformers model for DNA-language in genome. Bioinformatics, 37(15), 2112–2120.

[12] Zhou, Z., Ji, Y., Li, W., Dutta, P., Davuluri, R., and Liu, H. DNABERT-2: Efficient Foundation Model and Benchmark For Multi-Species Genome. (2023).

[13] Dalla-Torre, H., Gonzalez, L., Revilla, J. M., Carranza, N. L., Grzywaczewski, A. H., Oteri, F., Dallago, C., Trop, E., Sirelkhatim, H., Richard, G., Skwark, M., Beguir, K., Lopez, M., and Pierrot, T. (2023) The Nucleotide Transformer: Building and Evaluating Robust Foundation Models for Human Genomics. bioRxiv,.

[14] Nguyen, E., Poli, M., Faizi, M., Thomas, A., Birch-Sykes, C., Wornow, M., Patel, A., Rabideau, C., Massaroli, S., Bengio, Y., Ermon, S., Baccus, S. A., and Ré, C. HyenaDNA: Long-Range Genomic Sequence Modeling at Single Nucleotide Resolution. (2023).

[15] Benegas, G., Batra, S. S., and Song, Y. S. (2023) DNA language models are powerful predictors of genome-wide variant effects. Proceedings of the National Academy of Sciences, 120(44), e2311219120.

[16] Fishman, V., Kuratov, Y., Petrov, M., Shmelev, A., Shepelin, D., Chekanov, N., Kardymon, O., and Burtsev, M. (2023) GENA-LM: A Family of Open-Source Foundational DNA Language Models for Long Sequences. bioRxiv,.

[17] Mourad, R. (May, 2023) Semi-supervised learning improves regulatory sequence prediction with unlabeled sequences. BMC Bioinformatics, 24(1), 186.

[18] The ENCODE Consortium (September, 2012) An integrated encyclopedia of DNA elements in the human genome. Nature, 489(7414), 57–74.

[19] Zhang, Y., Liu, T., Meyer, C. A., Eeckhoute, J., Johnson, D. S., Bernstein, B. E., Nusbaum, C., Myers, R. M., Brown, M., Li, W., and Liu, X. S. (Sep, 2008) Model-based Analysis of ChIP-Seq (MACS). Genome Biology, 9(9), R137.

[20] Abramov, S., Boytsov, A., Bykova, D., Penzar, D. D., Yevshin, I., Kolmykov, S. K., Fridman, M. V., Favorov, A. V., Vorontsov, I. E., Baulin, E., Kolpakov, F., Makeev, V. J., and Kulakovskiy, I. V. (May, 2021) Landscape of allele-specific transcription factor binding in the human genome. Nature Communications, 12(1), 2751.

[21] Benegas, G., Batra, S. S., and Song, Y. S. (2022) DNA language models are powerful zero-shot predictors of non-coding variant effects. bioRxiv,.

[22] Xie, Q., Luong, M.-T., Hovy, E., and Le, Q. V. Self-training with Noisy Student improves ImageNet classification. (2020).

[23] Pham, H., Dai, Z., Xie, Q., Luong, M.-T., and Le, Q. V. Meta Pseudo Labels. (2021).

